# Phenotypic alteration of low-density granulocytes in people with pulmonary post-acute sequalae of SARS-CoV-2 infection

**DOI:** 10.1101/2022.10.22.513351

**Authors:** Logan S. Dean, Gehan Devendra, Boonyanudh Jiyarom, Natalie Subia, Michelle D. Tallquist, Vivek R. Nerurukar, Sandra P. Chang, Dominic C. Chow, Cecilia M. Shikuma, Juwon Park

## Abstract

Low-density granulocytes (LDGs) are a distinct subset of neutrophils whose increased abundance is associated with the severity of COVID-19. However, the long-term effects of severe acute respiratory syndrome coronavirus 2 (SARS-CoV-2) infection on LDG levels and phenotypic alteration remain unexplored. Using participants naïve to SARS-CoV-2 (NP), infected with SARS-CoV-2 with no residual symptoms (NRS), and infected with SARS-CoV-2 with chronic pulmonary symptoms (PPASC), we compared LDG levels and their phenotype by measuring the expression of markers for activation, maturation, and neutrophil extracellular trap (NET) formation using flow cytometry. The number of LDGs was significantly elevated in PPASC compared to NP. Individuals infected with SARS-CoV-2 (NRS and PPASC) demonstrated increased CD10^+^ and CD16HI subset counts of LDGs compared to NP group. Further characterization of LDGs demonstrated that LDGs from PPASC displayed higher NET forming ability and aggregation with platelets compared to LDGs from NP and NRS. Our data demonstrates that mature neutrophils with a heightened activation phenotype remain in circulation long after initial SARS-CoV-2 infection. Persistent elevation of markers for neutrophil activation and NET formation on LDGs, as well as an enhanced proclivity for platelet-neutrophil aggregation (PNA) formation in individuals with PPASC may be associated with the development of long-term pulmonary sequelae.

## 1 Introduction

Since December 2019, severe acute respiratory syndrome coronavirus 2 (SARS-CoV-2) has circulated worldwide, infecting over 600 million people globally while being responsible for over 6 million deaths (1). Although most people with COVID-19 infection recover within a few weeks, almost a third of survivors suffer from persistent symptoms, so called “Long COVID”, otherwise known as Post-Acute Sequalae of SARS-CoV-2 Infection (PASC) (2). PASC symptoms are highly variable and sustained for more than 30 days after initial COVID-19 infection (3). Development of Pulmonary PASC (PPASC), which includes persistent dyspnea and chronic cough, is not limited to recovered patients with prior hospitalizations or severe acute SARS-CoV-2 infection (4). Although studies have been initiated to identify factors associated with developing PPASC, mechanisms responsible for PPASC pathology remain unknown.

A growing body of literature implicates neutrophils as drivers of the hyperinflammatory state observed during acute COVID-19 (5–8). Increased neutrophil numbers are consistently reported during hospitalization due to COVID-19, and the neutrophil to lymphocyte ratio (NLR) is considered an independent risk factor for mortality among hospitalized COVID-19 patients (7). Moreover, Neutrophil Extracellular Traps (NETs), one of neutrophils’ defense mechanisms, is known to contribute to the maladaptive immune response via promotion of a hyperinflammatory state and via initiation of thrombotic events during COVID-19 (9–11). Histological examination of lung autopsies from severe COVID-19 patients reveal abundant NETs associated with microthrombi within alveolar capillaries (12). The combined evidence demonstrates that excessive neutrophils and NET production is associated with increased disease severity, pathophysiology, and poor clinical outcomes in COVID-19 (13).

Low-density granulocytes (LDGs) are a subpopulation of neutrophils that co-exist within peripheral blood mononuclear cell (PBMC) fractions (13). These cells demonstrate an enhanced ability to form NETs and are known to contribute to immune-mediated pathology through proinflammatory cytokine production and mediation of endothelial cell death (14–16). In the context of acute COVID-19 infection, frequencies of LDGs in individuals with mild to severe COVID-19 are increased compared to healthy controls (17,18). Additionally, their frequencies are positively correlated with markers for granulocyte recruitment and activation (19–22). Although elevated levels of LDGs and phenotypic activation are thought to play an important role in disease severity in COVID-19 patients, their circulating levels and functional contributions to long-term effects of COVID-19 and PPASC development are currently unknown.

In this study, we characterized LDG populations in individuals up to fourteen months following SARS-CoV-2 infection, alongside comparator groups, with and without, residual pulmonary symptoms. Our findings demonstrated that an altered LDG phenotype and increased NET and PNA formation are associated with prior SARS-CoV-2 infection and further enhanced in individuals with PPASC.

## 2 Methods

### Study Subjects & Specimens Selection

The “COVID-19 Infection in Hawaii” study is a cross sectional study of PASC complications in which participants were recruited from the state’s major tertiary care hospital. Participants completed written informed-consent documents and questionnaires regarding demographics, relevant medical history, and acute and/or lingering COVID-19 symptoms. PPASC patients were identified based on pulmonary function tests specifically a reduced predicted diffusing capacity for carbon monoxide (DLCOc, <80%) value, corrected for the patient’s hemoglobin. Comparitor study groups included patients fully recovered from SARS-CoV-2 infection with no residual symptoms >30 days after acute infection (NRS; n=10). Normal participants were selected from an existing HIV cohort study and who were HIV negative (NP; n=12). These groups were age- and gender-matched to the PPASC group for data analysis. The study was approved by the University of Hawaii at Manoa Human Studies Institutional Review Board (IRB#: RA-2020-053) and the Queens Medical Center Institutional Review Committee (IRB#: RA-2020-053).

### Plasma and PBMC isolation

Whole blood was collected in EDTA tubes by venipuncture and processed within 1 hour of collection based on previously published methods (23). Whole blood was centrifuged, plasma collected and cryopreserved at −80°C until downstream analysis was done. A separate aliquot of the whole blood was diluted with an equal volume of PBS overlayed onto Ficoll-Paque Plus (GE Healthcare Biosciences, Piscataway, NJ) according to manufacturer’s protocol. PBMC were aspirated off the Ficoll interphase, red blood cells lysed, and then washed twice in PBS supplemented with 1% Fetal Bovine Serum (FBS). Cells were then counted, viability determined, and cryopreserved at −80°C until further analysis.

### Quantification of cell-free DNA

Circulating cell-free DNA (cfDNA) in the plasma was quantified using the Quant-iT™ PicoGreen™ dsDNA Assay-Kit (ThermoScientific, P11496) according to the manufacturer’s instructions. Briefly, samples were diluted 1:20 in TE buffer and Quant-iT™ PicoGreen™ dsDNA reagent was added in a 1:1 ratio. After incubation for 5 min at RT in the dark, the fluorescence was read in a VICTOR3 plate reader (Perkin Elmer). The concentration of the circulating cell-free DNA was calculated using the standard provided by the kit.

### Enzyme-linked immunosorbent assay (ELISA) for NETs

96-well Enzyme ImmunoAssay/Radio ImmunoAssay (EIA/RIA) plates (#3590, Costar) were coated overnight at 4°C with an anti-elastase antibody (1:250, #sc-9521, Santa Cruz Biotechnology) in 15 mM of Na_2_CO_3_, 35 mM of NaHCO_3_, at pH 9.6. The next day, the wells were washed three times with PBS, blocked in 5% BSA for two hours at RT, and washed three times with PBS. Then, 50 μl of plasma samples were added to the wells, incubated for two hours at room temperature on a shaker, and then washed three times with wash buffer (1% BSA, 0.05% Tween 20 in PBS). Next, anti-DNA-peroxidase conjugated antibody (1:50, #11774425001, Roche) in 1% BSA in PBS was added to the wells for 2 hours at room temperature, and the wells were washed five times with wash buffer before the addition of 2,2’-azino-bis (3-ethylbenzothiazoline-6-sulphonic acid (ABTS, #37615, Thermo Fisher Scientific). Optical density was read 40 min later at 405 nm using a plate reader (SpectraMax i3, Molecular Devices).

### Flow Cytometric Analysis

Typically, 1-2×10^6^ PBMC were incubated with Fixable Viability Dye eFluor 506 (Ebiosciences, 1:1000) at 4°C for 30 minutes, followed by addition of Human TruStain FcX (BioLegend, San Diego, CA, 1:200) in flow buffer (HBSS supplemented with 1% BSA) at room temperature (RT) for 15 minutes. Subsequently, cells were stained with the titrated fluorophore-conjugated primary antibody cocktail (**S1 Table**) at RT for 30 minutes and then washed twice with ice-cold flow buffer. For the intracellular staining, cells were resuspended in 250 mL of BD Cytofix/Cytoperm (BD Technologies, East Rutherford, NJ) for 30 minutes at 4°C and then incubated with the titrated primary anti-MPO-PE and primary citrullinated histone H3 (citH3) antibody (Abcam, Waltham, MA) for 30 minutes at 4° C, protected from light. Allophycocyanin (APC)-conjugated anti-goat secondary antibody (Invitrogen, Waltham, MA) at 1:500 dilution was added for labeling the anti-citH3 primary antibody. Samples were then washed twice with ice-cold flow buffer and resuspended to 800 mL of flow buffer for acquisition. Samples were acquired on a Attune NxT Flow Cytometer (Thermofisher, Waltham, MA) with approximately 1.0×10^6^ events collected per sample. Data was normalized to the amount of acquired events and analyzed using FlowJo (Treestar, Ashland, OR) software.

### Statistics

This is a cross sectional study comparing PPASC, NRS, and NP participants. Flow cytometry results were analyzed using Mann Whitney-U test with a 95% confidence interval for between groups comparisons. Spearman correlations were performed for between variable associations with a linear regression model for multi-variate analysis. A p value <0.05 was considered statistically significant for all tests. Statistical analysis was performed using Prism 9 (Graph Pad, San Diego, CA).

## 3 Results

### Circulating LDGs display increased frequency of maturation/activation

To understand the impact of long-term COVID-19 on LDGs and LDG dysregulation in pulmonary PASC, we selected samples from COVID-19 convalescents with pulmonary symptoms (PPASC group [PPASC], n=12), COVID-19 convalescents with no residual symptoms (recovered group [NRS], n=10). A comparator group of HIV-seronegative controls from a previous HIV cohort study was used as a SARS-CoV-2 naïve control (naïve group [NP], n=12). All groups were matched for age and gender. The baseline participant characteristics are displayed in **Table 1**. The median age of participants was 57, 55, and 54.5 years for NP, NRS, and PPASC participants, respectively. The majority of participants were male (84.4% of all participants) and the prevalence of pre-existing conditions, except for diabetes, was similar between groups. PPASC participants had a significant increase in BMI compared to NP, but not NRS.

**Table 1:** Demographics and Pre-existing Conditions of naive participants (NP), no residual symptoms (NRS), and pulmonary PASC (PPASC) participants.

|  |  |  |  |  |
| --- | --- | --- | --- | --- |
|  |  | **NP (n=12)** | **NRS (n=10)** | **PPASC (n=12)** |
| **Demographics** | Age (years) | 57.0 [53.3-63.5] | 55.0 [43.0-60.0] | 54.5 [49.8-58.0] |
|  | Male (n, %) | 10 (83.3) | 7 (70.0) | 10 (83.3) |
|  | Caucasian (n, %) | 9 (75.0) | 3 (30.0) | 4 (33.3) |
|  | BMI (kg/m<sup>2</sup>) | 27.8 [23.9-29.0] | 26.3 [20.6-33.0] | 30.8 [26.2-34.1]* |
|  | Time since Infection (months) | 0 [0] | 5 [1.5-9.5] | 6 [4-14] |
| **Pre-Existing Conditions** | Diabetes (n, %) | 1 (8.3) | 1 (10.0) | 4 (33.3) |
|  | Hypertension (n, %) | 4 (33.3) | 2 (20.0) | 5 (41.7) |
|  | History of Smoking (n, %) | 6 (50.0) | 3 (30.0) | 4 (33.3) |
|  | COPD (n, %) | 0 (0) | 0 (0) | 0 (0) |
|  | Asthma (n, %) | 3 (25.0) | 1 (10.0) | 2 (16.7) |

Values presented as [median, Q1-Q3] unless otherwise stated. \*p<.05 compared to NP.

LDGs were identified as CD45^+^/CD11b^+^/CD14^-^/CD16^+^/CD15^+^ cells (**Figure 1A**). The PPASC group demonstrated an increasing trend in the percentage of LDGs (1.1% [0.40-3.50%]) (**Figure 1B**) and significantly elevated LDG counts (1243 [358-2421]) as compared to NP (0.67% [0.45-1.14%]; 372 [95-652], respectively) (**Figure 1C**). Interestingly, LDG counts were elevated, albeit not significantly, in NRS than NP although their frequencies were comparable to NP, suggesting that there is preferential expansion of LDG populations in NRS.

**Figure 1.**
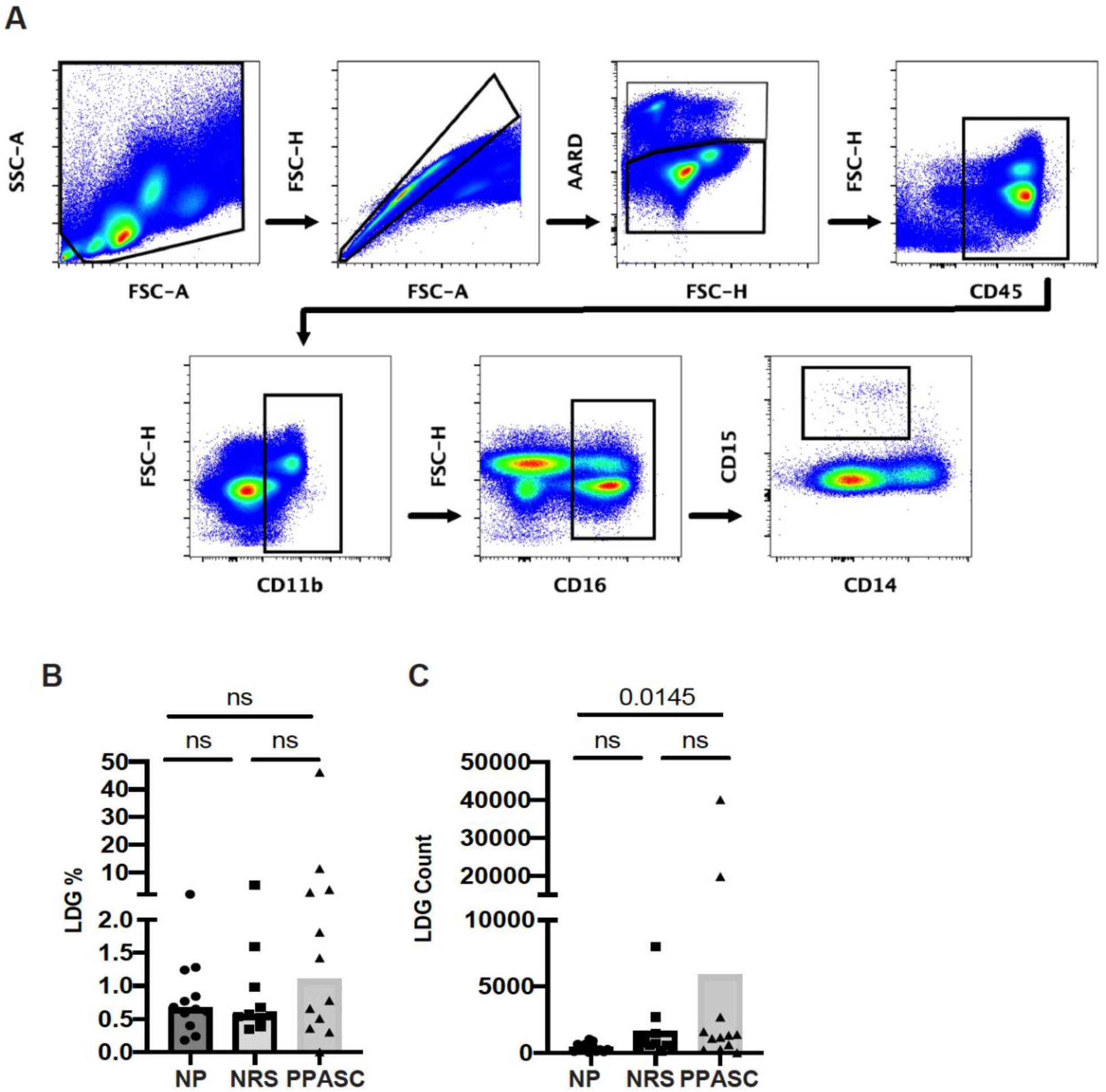
Circulating low-density granulocytes (LDGs) flow cytometry gating strategy. Representative gating for LDG identification and LDGs were identified by CD15 expression after gating on CD45^+^CD11b^+^CD16^+^CD14^-^ cells. Box region within scatter plot represents LDGs within NP (n=12), NRS (n=10), and PPASC (n=12) groups (**A**). Percentage of LDG (**B**) and total count of LDGs in NP, RC, and PPASC groups (**C**). Graphs are shown as bar with scatter dot plots. Comparisons between groups were performed using Mann-Whitney U test. ns; non-significant, p >0.05.

Frequencies of LDGs positive for CD10, known as a mature granulocyte marker, were significantly increased in NRS and PPASC as compared to NP (**Figure 2A**). CD10^+^ LDG counts also were significantly increased in PPASC (p=0.007) and NRS (p=0.014) compared to NP (684 [182-1766], 337 [190-913], 105 [31-243], respectively) (**Figure 2B**).

**Figure 2.**
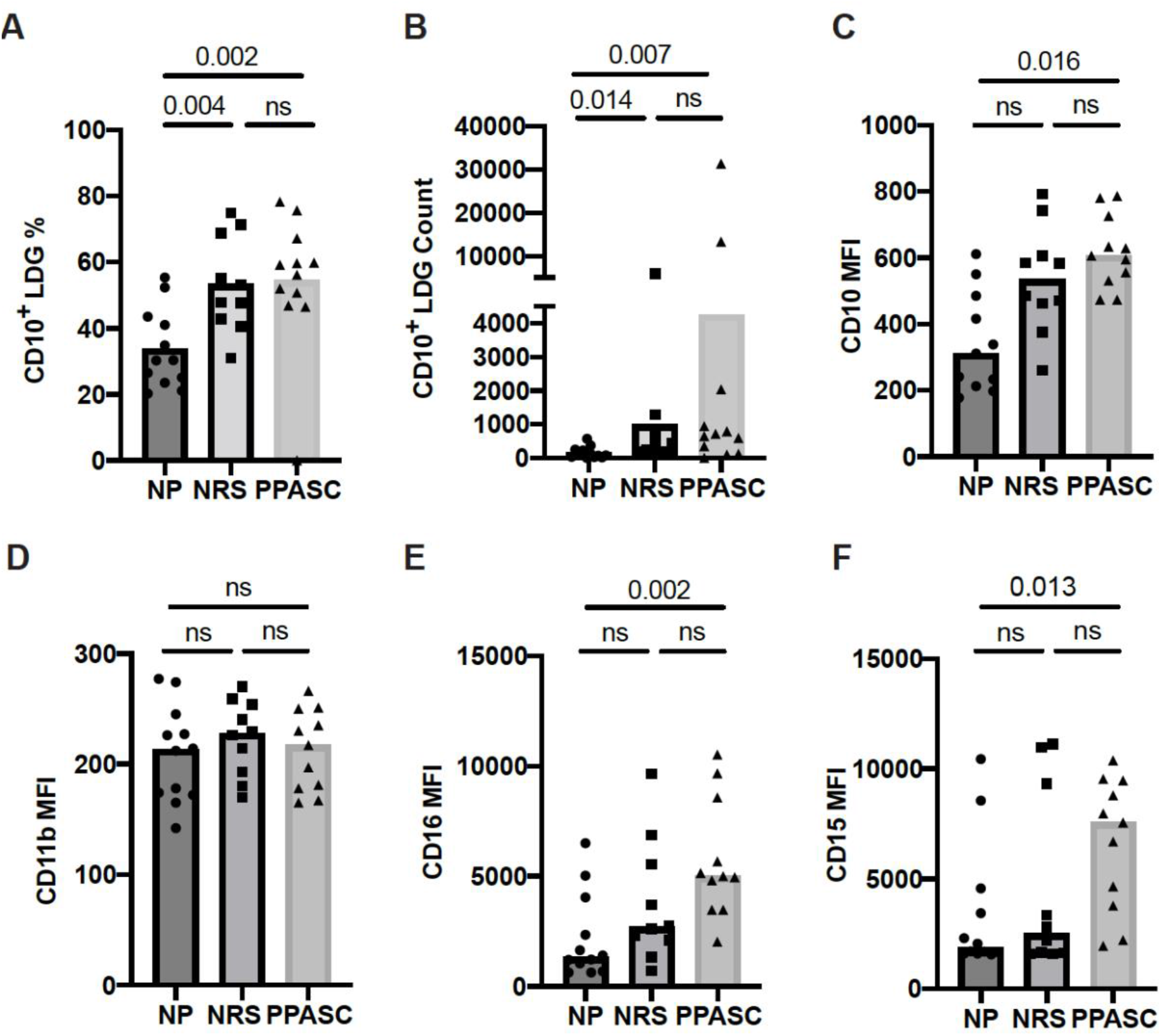
LDG display a mature phenotype in COVID-19 convalescents. Mature LDGs were identified by their CD10 expression. The percentage (**A**) and total number (**B**) of CD10^+^ cells in the LDG fraction were compared across three groups. Analysis of surface marker expression on circulating LDGs and median fluorescence intensity (MFI) of CD10 (**C**), CD11b (**D**), CD16 (**E**), and CD15 (**F**) on LDGs in NP, NRS, and PPASC groups. Graphs are shown as bar with scatter dot plots. Comparisons between groups were performed using Mann-Whitney U test. ns; non-significant, p > 0.05.

Both LDGs from NRS and PPASC displayed higher median fluorescence intensity (MFI) of CD10 than NP (**Figure 2C**). Indeed, MFI expression of CD15 and C16 per granulocyte was significantly increased in PPASC compared to NP, although there was a rising trend of increased expression within NRS (**Figure 2E, F**). Next, we assessed the activation status of LDGs by determining CD11b and MPO expression and granularity. There was no difference in MFI expression of CD11b and granularity between the three groups (**Figure 2D and 3**). Taken together, our data indicate that circulating LDGs display increased frequency of maturation and their levels remain elevated for many months after infection, even within patients who recovered from COVID-19 with no residual symptoms.

**Figure 3.**
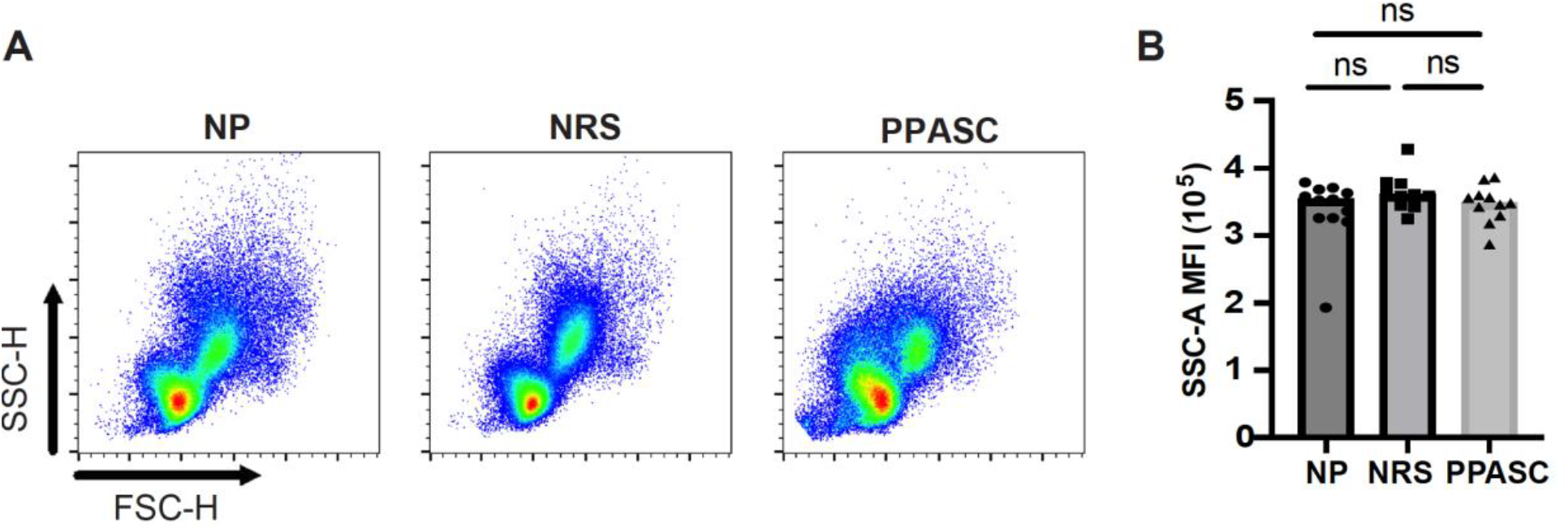
Granularity of LDGs after SARS-Cov-2 Infection. Morphology (size and granularity) of LDGs were analyzed by flow cytometry. Gating strategy for identification of granularity on LDG after gating on CD45^+^CD11b^+^CD16^+^CD14^-^CD15^+^cells (**A**). Mean granularity of LDGs based on sideward scatter/area of intensity (SSC-A) in NP, NRS, and PPASC (**B**). Graphs are shown as bar with scatter dot plots. Comparisons between groups were performed using Mann-Whitney U test. ns; non-significant, p > 0.05.

### PPASC exhibits elevated levels of CD16^hi^ LDGs in circulation

Recent studies have identified different LDG subsets based on the relative expression levels of CD16 in COVID-19 patients (24–27). Our data indicated that LDG from NRS and PPASC exhibited more mature neutrophils, compared to LDG from NP. Indeed, the trend of higher CD16 MFI expression was detected on LDG from PPASC, which led us to further examine if a specific LDG subset (based on CD16 expression) is increased in PPASC. Similar to previous observations, we found two LDG subsets (CD16^lo^ and CD16^hi^) within PBMC (**Figure 4A**). CD16^hi^ LDG from PPASC displayed a mature neutrophil phenotype denoted by CD10 expression (**Figure 4B**). Notably, increased frequencies and numbers of CD16^hi^ LDG appeared in PPASC, compared with NRS and NP (**Figure 4A, C-F**). These data suggest that higher CD16 MFI expression in PPASC is related to expansion of CD16^hi^ LDG.

**Figure 4:**
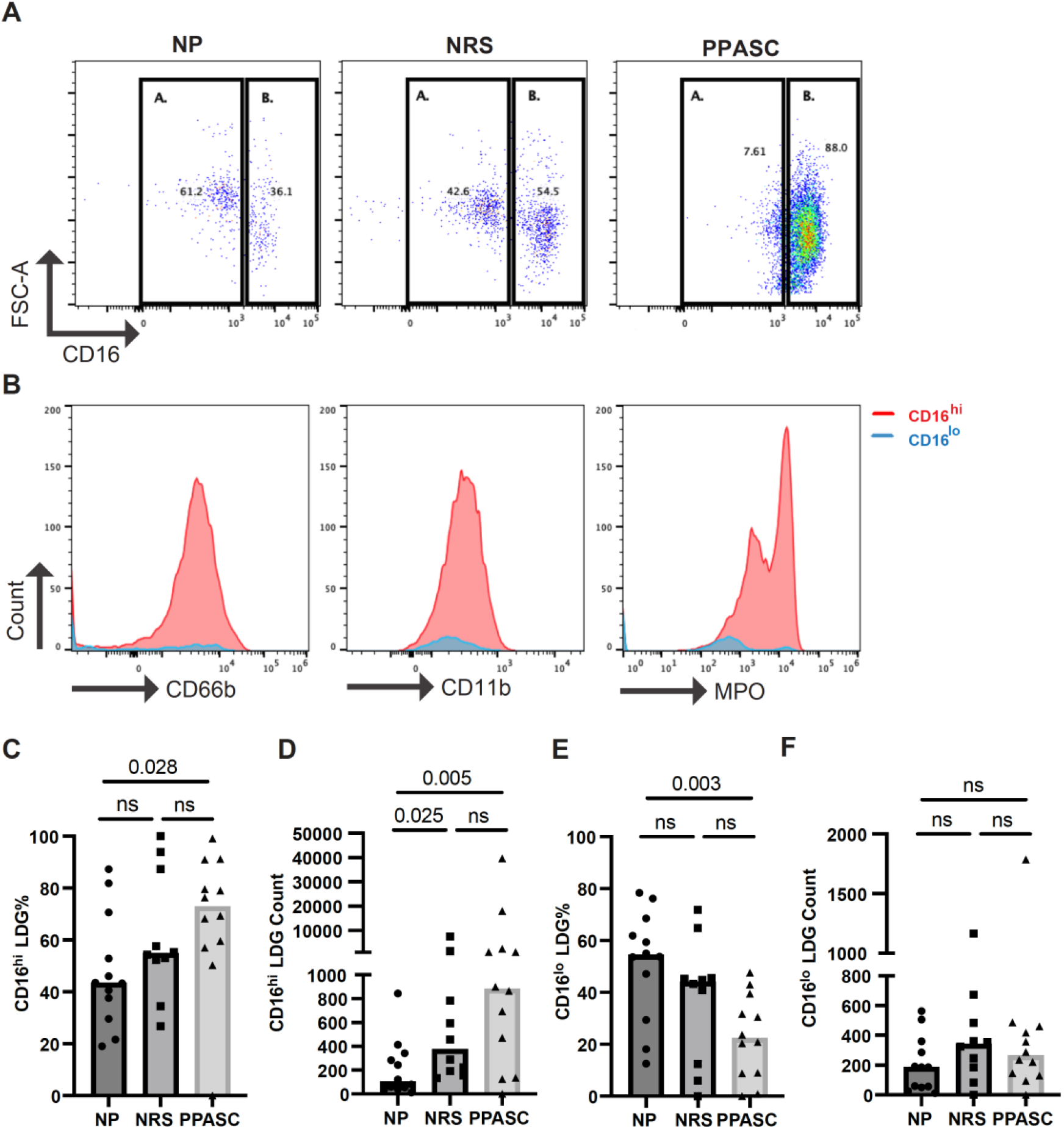
Representative dot plot showing LDG populations in NP, NRS, and PPASC separated into CD16^lo^ (A) and CD16^hi^ (B) populations (**A**) Representative histograms showing expression of CD66b, CD11b, and MPO in LDG populations from PPASC participants according to CD16^lo^ (blue) and CD16^hi^ (red) delineation (**B**) CD16^hi^ LDG % in NP, NRS, and PPASC groups (**C**) CD16^hi^ LDG counts in NP, NRS, and PPASC groups (**D**) CD16^lo^ LDG % in NP, NRS, and PPASC groups (**E**) CD16^lo^ LDG count in NP, NRS, and PPASC groups (**F**) Mann-Whitney U Test, ns; non-significant, p >0.05

### LDGs from PPASC demonstrate an enhanced ability to form NETs

COVID-19 convalescents showed increased LDGs displaying mature phenotype. Indeed, MFI of MPO expression on LDGs was higher in these individuals (**Figure 5D**), suggesting that COVID-19 convalescents may possess more abundant granules within the neutrophils. LDGs have been proposed as the main source of NETs due to their enhanced capacity of spontaneous NET release in autoimmune and inflammatory diseases. We further investigated if LDGs from PPASC appear more prone to form NETs. NET forming LDGs were identified as double positive for citrullinated histone H3 (CitH3) and MPO by flow cytometry (**Figure 5A**). Although the frequency of LDGs able to form NETs was decreased in PPASC, the numbers of NET forming LDGs were significantly increased in both NRS and PPASC, compared to NP (**Figure 5B, C**). In addition, MPO expression levels derived from NET forming LDGs in PPASC were significantly increased compared to NP, potentially indicating increased neutrophil activation in PPASC (**Figure 5E**). To verify whether plasma of COVID-19 convalescents contained increased concentrations of NETs as a consequence of increased numbers of circulating LDGs with enhanced ability to release NETs, we quantified plasma level of cell-free DNA (cfDNA) and CitH3. As shown in Figure 5F and G, we observed elevated cfDNA in PPASC compared to NP, and elevated H3cit levels in NRS and PPASC, compared to NP. These results are the first to demonstrate an increase in NET forming LDGs and plasma NET levels in convalescent SARS-CoV-2 infected individuals following COVID-19.

**Figure 5.**
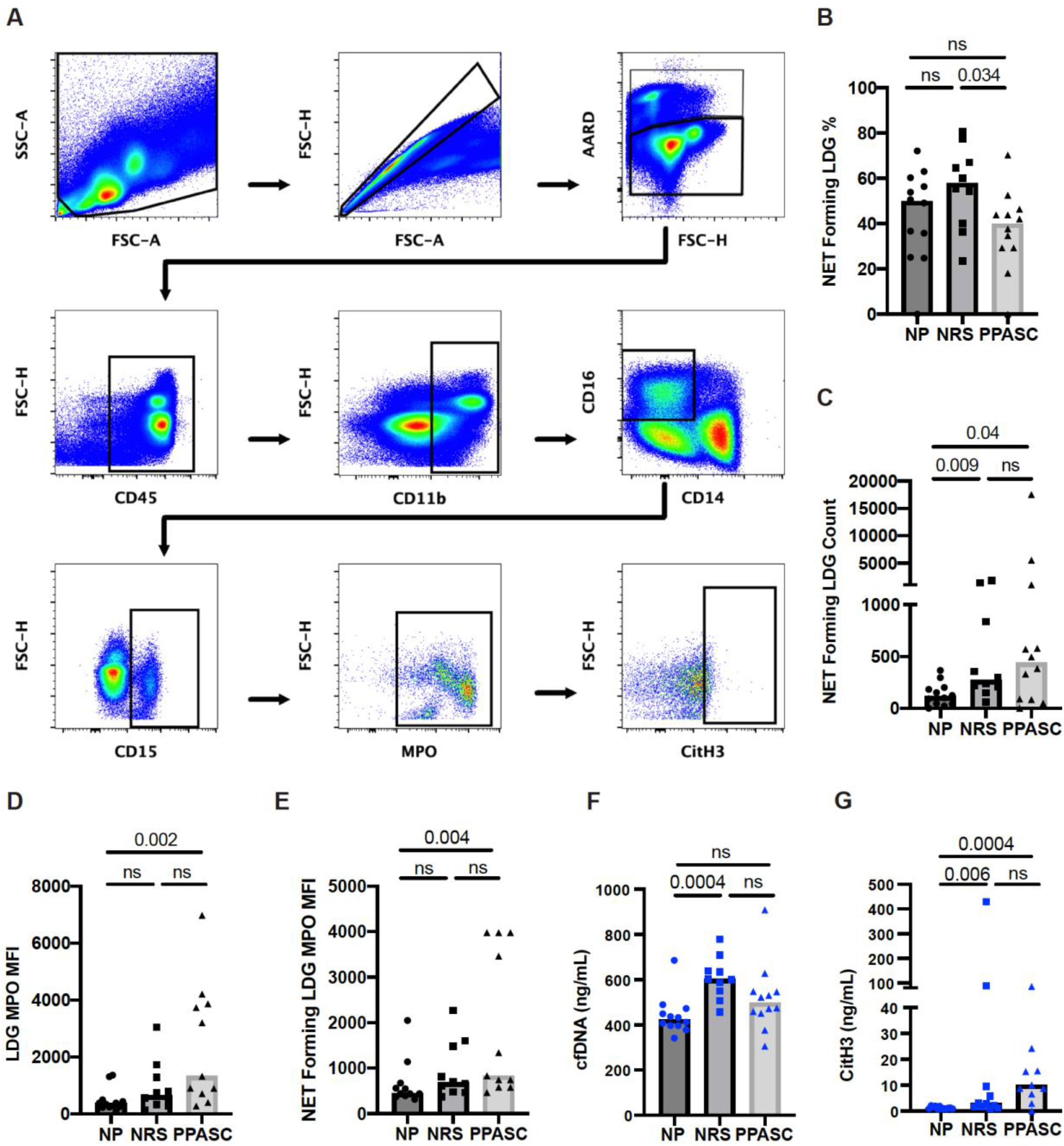
Characterization of NET formation in LDGs. Flow cytometric determination of NET forming LDGs. Whole PBMC were gated on side scatter area (SSC-A) and forward scatter area (FSC-A) to gate out debris. Forward scatter height (FSC-H) and FSC-A gated out doublets followed by a viability gate using LIVE/DEAD stain and FSC-A. Leukocytes were selected by CD45 expression and CD11b^+^CD14^-^CD16^+^ fractions were finally identified for LDG based on CD15 expression. NETs were defined as MPO and CitH3 double positive events within the LDG population (**A**). The percentage (**A**) and total number (**B**) of NET forming cells in the LDG fraction was compared across three groups. MFI of MPO on LDGs (**C**) and NET forming LDGs (**D**) in NP, NRS, and PPASC groups. Graphs are shown as bar with scatter dot plots. Comparisons between groups were performed using Mann-Whitney U test. ns; non-significant, p > 0.05.

### Platelets form aggregates with LDGs in COVID-19 infected individuals

After observing increased NET forming LDGs in both PPASC and NRS individuals, we hypothesized that an increase in NETs would favor PNA. Although the number of platelets was no different between three groups (**Figure 6A**), we observed a significant increase in the numbers of platelets bound to LDGs within the PPASC group (**Figure 6B**). As NETs have been demonstrated to provide scaffolding for platelet adhesion, activation, and aggregation, we examined platelet populations bound specifically to NET forming LDG populations. As expected, COVID-19 convalescents showed an increase in platelet binding to NET forming LDGs, which is more prominent in COVID-19 convalescents who had residual pulmonary symptoms (**Figure 6C**). To determine if activated platelet facilitate PNA in COVID-19, we examined CD62P (P-selectin) expression on platelets bound to NET forming LDGs. MFI expression of P-selectin was not different between three groups, suggesting that PNA is independent of P-selectin expression on platelets (**Figure 6D**).

**Figure 6.**
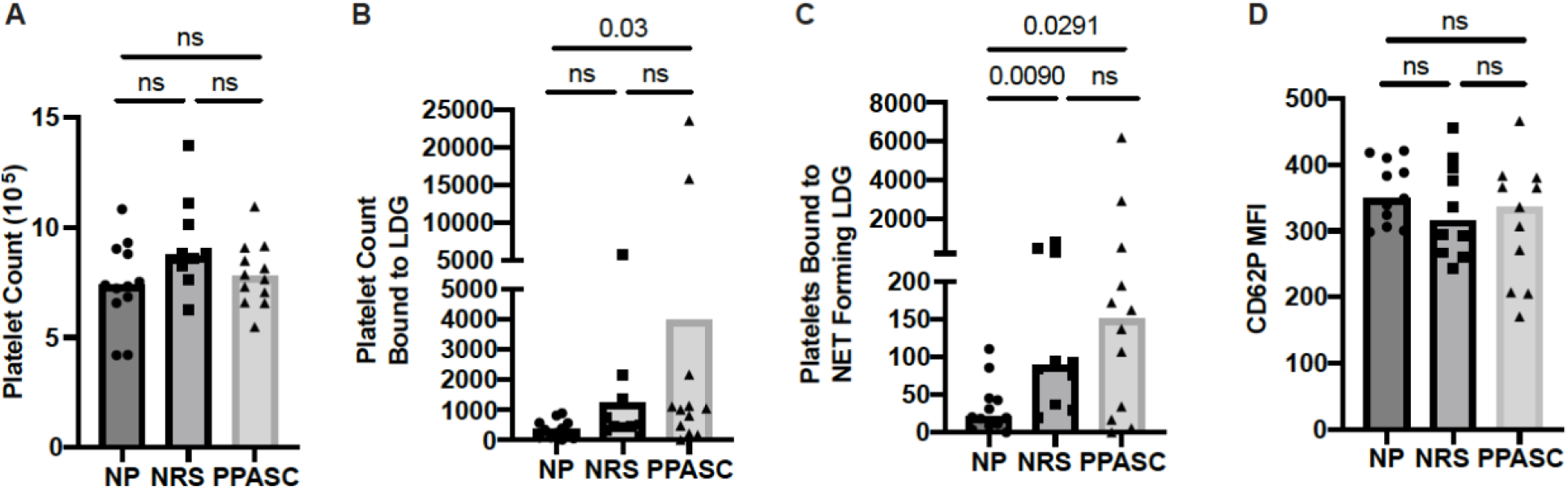
Total number of platelets (**A**), bound to LDGs **(B)**, bound to NET forming LDGs **(C)**, and MFI of CD62P on platelet bound to NET forming LDGs in NP, NRS, and PPASC (**D**). Graphs are shown as bar with scatter dot plots. Comparisons between groups were performed using Mann-Whitney U test. ns; non-significant, p > 0.05

## 4 Discussion

A comprehensive understanding of the host immune response and the immunopathological characteristics in COVID-19 individuals with PASC is important for developing therapeutic treatments to target lingering symptoms. Elevated levels of LDG and their phenotypic activation are thought to play an important role in disease severity in COVID-19 patients (28). However, their concentration in circulation and functional contributions to long-term effects of COVID-19 are currently unknown. In this study, we analyzed a cross-sectional LDG population and compared circulating LDG levels in patients with PPASC and with no residual symptoms to individuals naïve to SARS-CoV-2. We also investigated the phenotypic and functional alteration of LDGs in the study participants. To the best of our knowledge, there is currently no report regarding the long-term effect of SARS-CoV-2 on LDGs in COVID-19 individuals and this is the first time it has been shown that circulating LDGs remain elevated during PPASC onset, following SARS-CoV-2 infection.

Even more remarkable was the observation that profound increases in CD10^+^, mature LDG numbers were observed in both NRS and PPASC groups. Specifically, the increased CD10^+^ LDG counts in these individuals were reflected as a significant increase in the proportion of CD10^+^ cells within the LDG population. Previous studies have highlighted that increased numbers of immature granulocytes in the blood of patients with severe COVID-19 correlates with the disease severity (29) Torres-Ruiz et al. (2021) reported that both CD10^-^ (immature) and CD10^+^ (mature) LDG population were significantly higher in severe/critical COVID-19 patients. In context of COVID-19, LDGs expressing CD10^low^ resemble myeloid-derived suppressor cells (30,31), in contrast mature LDGs with higher CD10 and CD16 expression are more prone to form NETs (32). In line with previous findings (33), LDGs from COVID-19 convalescents demonstrated more mature granulocytes with higher expression of CD10, CD16, and MPO, in combination with enhanced NET production.

We futher found that the proportions of CD16^hi^ LDGs were significantly higher in PPASC than NP, in contrast to the reduced proportions of CD16^lo^ LDGs in the PPASC group. As previously described, neutrophil morphology in COVID-19 patients vary between mature segmented and immature round nuclei (34). Moreover, a majority of CD16^int^ neutrophils contained band-shaped nucleus, but CD16^hi^ neutrophils were bilobed rather than hypersegmented, a similar morphology observed in other severe infections (35). These findings suggest the possibility of morphological abnormalities in COVID-19 individuals even in mature neturophils. Although morphological analysis of neutrophils was not performed in this study, CD16^hi^ LDGs showed enhanced neutrophil activation and displayed higher MFI of CD66b, CD11b, and MPO. Our findings demonstrate that the activated function of mature LDGs and their high affinity for platelet aggregation are associated with lung inflammation and thrombosis among individuals with PPASC. The maturation status of LDG and their imparied functionality during long-term symptoms associated with post-SARS-CoV-2 infection should be of prominent interest for understanding the association of LDGs with PPASC onset and persistence.

Our study was limited in multiple ways. Our sample size was small. The length of time following COVID-19 confirmation and enrollment was variable, ranging from 1 to 10 months post-infection. This variability in time of sample collection may have influenced LDG population characteristics. Thus, a longitudinal evaluation of neutrophil dynamics and functional properties after COVID-19 infection should be carried out to determine whether neutrophil dysfunction is associated with the clinical outcome of respiratory failure. Hematological analysis would provides direct evidence of changes in whole blood cellular composition, however complete blood counts (CBC) were not performed on study participants. As a result, a comparison of our flow cytometric evaluations to clinical CBC counts was not possible.

In conclusion, our data demonstrate that SARS-CoV-2 infection leads to long-lasting alterations in the peripheral LDG functionality and activation, including enhanced NET formation and platelet aggregation. These alterations may contribute to increased tissue damage and thrombosis following infection with SARS-CoV-2 and suggest that sustained expansion of neutrophils subsets may correlate with prolonged respiratory symptoms. Further studies are needed to determine whether targeting of the neutrophil subset, including LDGs, would be beneficial to prevent the development and persistence of PPASC.

## Supporting information

Antibody-fluorophore conjugations and company used for flow cytometry

