## Supplementary material for "Phenotypic alteration of low-density granulocytes in people with pulmonary post-acute sequalae of SARS-CoV-2 infection": Antibody-fluorophore conjugations and company used for flow cytometry

**Supplementary Table 1**

| **Antibody Target** | **Conjugate Fluorophore** | **Company** |
| --- | --- | --- |
| CD45 | BV711 | BD Biosciences (East Rutherford, NJ) |
| CD11b | PE-Cy-7 | BioLegend (San Diego, CA) |
| CD14 | BV605 | BioLegend (San Diego, CA) |
| CD16 | BV650 | BioLegend (San Diego, CA) |
| CD15 | FITC | Millipore Sigma (St. Louis, MO) |
| CD10 | PerCP5.5 | R&D Systems (Minneapolis, MN) |
| CD41 | PE-Dazzle | BioLegend (San Diego, CA) |
| CD62p | AF700 | BioLegend (San Diego, CA) |
| CD66b | BV421 | BioLegend (San Diego, CA) |
| MPO | PE | BD Biosciences (East Rutherford, NJ) |
| citH3 | APC | Abcam & Invitrogen (Waltham, MA), respectively |
| Viability | eFluor506 | Invitrogen (Waltham, MA) |

**Supplementary Table 1.** Antibody-fluorophore conjugations and company used for flow cytometry
